# Structural basis for autophagy inhibition by the human Rubicon-Rab7 complex

**DOI:** 10.1101/2020.04.18.048462

**Authors:** Hersh K. Bhargava, Keisuke Tabata, Jordan M. Byck, Maho Hamasaki, Daniel P. Farrell, Ivan Anishchenko, Frank DiMaio, Young Jun Im, Tamotsu Yoshimori, James H. Hurley

## Abstract

Rubicon is a potent negative regulator of autophagy and a potential target for autophagy-inducing therapeutics. Rubicon-mediated inhibition of autophagy requires the interaction of the C-terminal Rubicon homology (RH) domain of Rubicon with Rab7-GTP. Here we report the 2.8 Å crystal structure of the Rubicon RH domain in complex with Rab7-GTP. Our structure reveals a novel fold for the RH domain built around four zinc clusters. The switch regions of Rab7 insert into pockets on the surface of the RH domain in a mode that is distinct from those of other Rab-effector complexes. Rubicon residues at the dimer interface are required for Rubicon and Rab7 to colocalize in living cells. Mutation of Rubicon RH residues in the Rab7 binding site restore efficient autophagic flux in the presence of overexpressed Rubicon, validating the Rubicon RH domain as a promising therapeutic target.

## Introduction

Macroautophagy (hereafter, “autophagy”) degrades bulky cytoplasmic materials by encapsulating them in a double membrane structure, the autophagosome, and delivering them to the lysosome for degradation^1^. Autophagy is critical for maintaining cellular homeostasis, and functions as a response to events such as starvation, pathogenic invasion, damage to organelles, and toxic protein aggregation^2^. Failure of autophagy is associated with aging, cancer, heart disease, neurodegeneration, and metabolic disorders^3^. Autophagy regulators are therefore key targets for the development of novel therapeutics^4, 5^. Drugs that directly upregulate autophagy, whether by activation of a positive regulator or inhibition of a negative regulator, are highly sought after.

Rubicon (Run domain Beclin-1 interacting and cysteine-rich containing) is a potent negative regulator of autophagy and endolysosomal trafficking^6, 7^. Rubicon disrupts autophagosome-lysosome fusion by inhibiting the class III phosphatidylinositol 3-kinase complex 2 (PI3KC3-C2)^6–8^. Rubicon is targeted to its sites of action by the late endosomal small GTPase Rab7^9, 10^. Autophagy suppression by Rubicon in aging has been linked to decreased clearance of α-synuclein aggregates in neural tissues, impairment of liver cell homeostasis, and interstitial fibrosis in the kidney^11–13^. Together with another target for autophagy inducing therapeutics, Bcl-2^14, 15^, Rubicon is one of very few known broadly-acting negative regulators of autophagy. Consistent with the concept that decreased autophagic function is associated with aging, Rubicon knockout increases lifespan in worms and flies^12^. Most encouragingly, Rubicon knockout mice show decreases in multiple age-associated pathologies, including kidney fibrosis and α-synuclein accumulation^12^.

Rubicon consists of an N-terminal RUN domain with unknown function, a middle region (MR) which contains its PI3K binding domain (PIKBD)^8^, and a C-terminal Rubicon homology (RH) domain which mediates interaction with Rab7^9^ (Fig. 1a). Rab7 is the characteristic Rab marker of late endosomes^16^. Rubicon is the defining member of a class of metazoan RH domain-containing autophagy regulators, which also includes PLEKHM1 and Pacer^9, 17^, two other proteins that modulate late steps in autophagy. The RH domains of Rubicon, PLEKHM1, and Pacer all contain nine conserved cysteines and one conserved histidine, which have been predicted to bind divalent zinc cations and are required for Rubicon and PLEKHM1 to interact with Rab7^9^. Despite the importance of the RH domain in targeting Rubicon and other regulators of the late stages of autophagy, no structural information has been available for the RH domain of any of these proteins.

**Figure 1:**
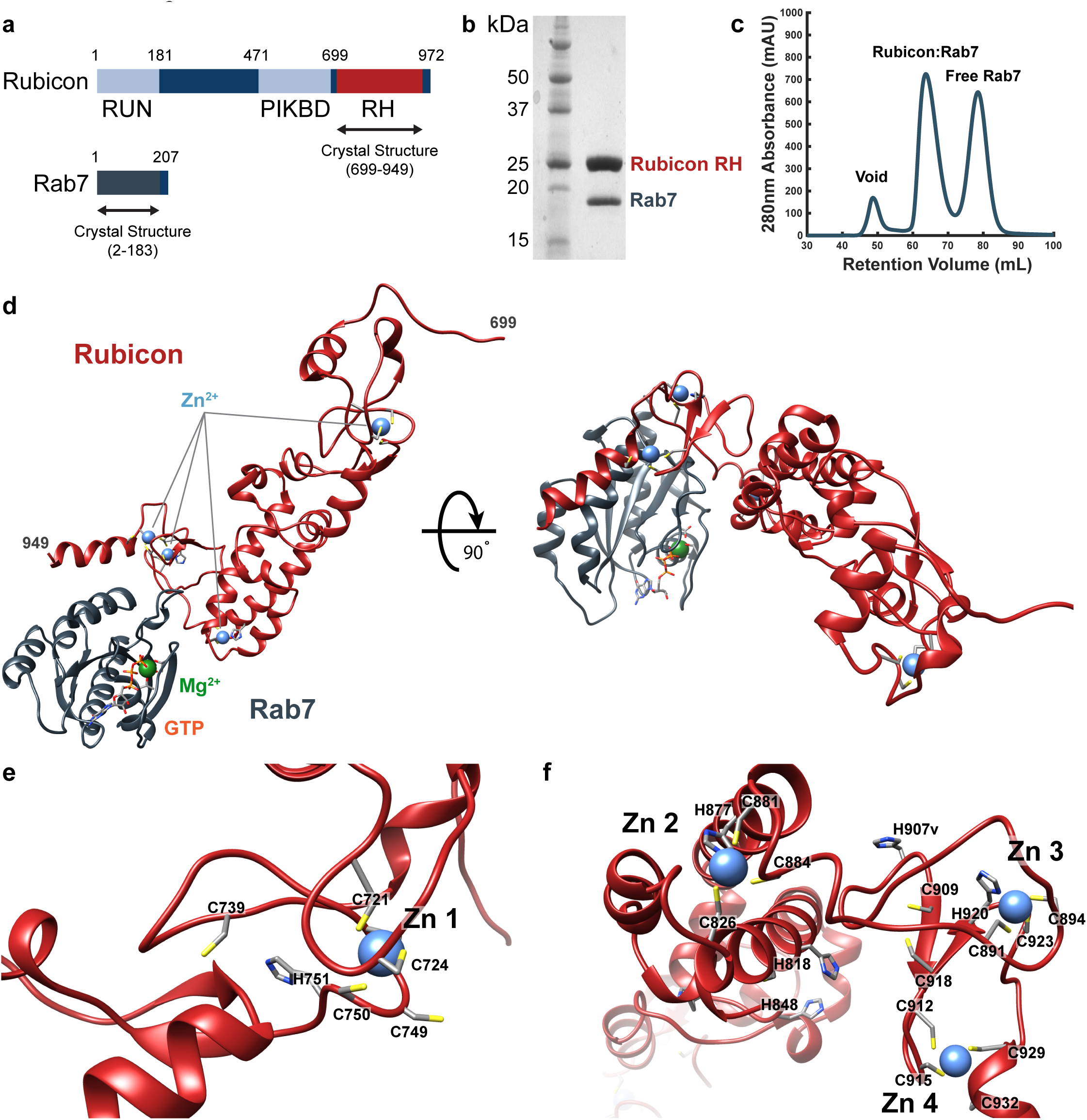
Structure of the Rubicon RH domain in complex with GTP-bound Rab7. (a) Schematic representation of the domain organizations of human Rubicon and Rab7. The regions crystallized are indicated with black arrows. (b) SDS-PAGE analysis of the Rubicon:Rab7 sample yielding the crystal used for structure determination. (c) Size exclusion chromatography (SEC) chromatogram of the mixture of Rubicon RH and Rab7 in a 1:1.5 molar ratio. (d) Overall structure of the human Rubicon RH domain in complex with GTP-bound Rab7. Zinc, magnesium, and GTP ligands are annotated. (e) Structure of the first zinc finger motif of Rubicon and nearby cysteine and histidine residues. (f) Structure of zinc finger motifs 2, 3, and 4 and nearby cysteine and histidine residues.

We sought to determine the structural basis for Rab7-dependent recruitment of Rubicon with the long-term goal of blocking this step therapeutically. The structure revealed a novel, four zinc-cluster based fold for the Rubicon RH domain, and revealed how Rubicon and Rab7 interact. Rubicon mutants that disrupted the Rubicon:Rab7 interaction were shown to mislocalize and to be incapable of suppressing autophagy, validating this interface as a potential drug target.

## Results

### Structure of the Rubicon RH domain

We expressed a soluble 29.4 kDa Rubicon RH domain construct comprising Rubicon residues 699-949. It was essential to supplement the expression medium with ZnCl_2_ to obtain measurable quantities of Rubicon. Rubicon RH bound stably to Rab7-GTP in size exclusion chromatography (Fig. 1b-c), where Rab7-GTP refers to a GTP-locked Q67L mutant construct comprising residues 2-183. The C-terminal tail of Rab7, which ends in a dicysteine motif that is prenylated *in vivo*, was truncated in this construct. We determined the structure of the Rubicon RH-Rab7-GTP complex (Fig. 1d, 2a) by x-ray crystallography at a resolution of 2.8 Å. Phases were determined by molecular replacement with a search model based on contact prediction from co-evolutionary data^18^. The asymmetric unit of the crystal contains a single Rubicon RH-Rab7 complex. Statistics of the crystallographic data collection and structure refinement are provided in Table 1, and representative electron density is displayed in Fig. 2b.

**Table 1:** Crystallographic Data Collection and Refinement Statistics

| Data Collection |  |
| --- | --- |
| Space Group | *I*2<sub>1</sub> |
| Unit Cell Dimensions<br>a, b, c (Å)<br>$\alpha$, $\beta$, $\gamma$ (°) | 60.04, 45.33, 186.28<br>90.00, 98.51, 90.00 |
| Resolution (Å) | 92.11 - 2.80 (2.95 - 2.80) |
| R<sub>pim</sub> | 0.1638 (0.3938) |
| I/$\sigma$(I) | 6.4 (2.2) |
| Completeness (%) | 99.1 (98.8) |
| Redundancy | 2.8 (2.8) |
| Refinement |  |
| Resolution (Å) | 92.11 - 2.80 |
| No. of reflections | 171,883 (17,031) |
| R<sub>work</sub>/R<sub>free</sub> (%) | 19.39/26.54 |
| Number of atoms<br>Protein<br>Ligand/Ion | 3414<br>33 |
| B factors protein (Å<sup>2</sup>)<br>Protein<br>Ligand/Ion | 58.06<br>49.57 |
| RMS Deviations<br>Bond Lengths (Å)<br>Bond Angles (°) | 0.009<br>1.05 |
| Values in parentheses are for the highest-resolution shell. |  |

**Figure 2:**
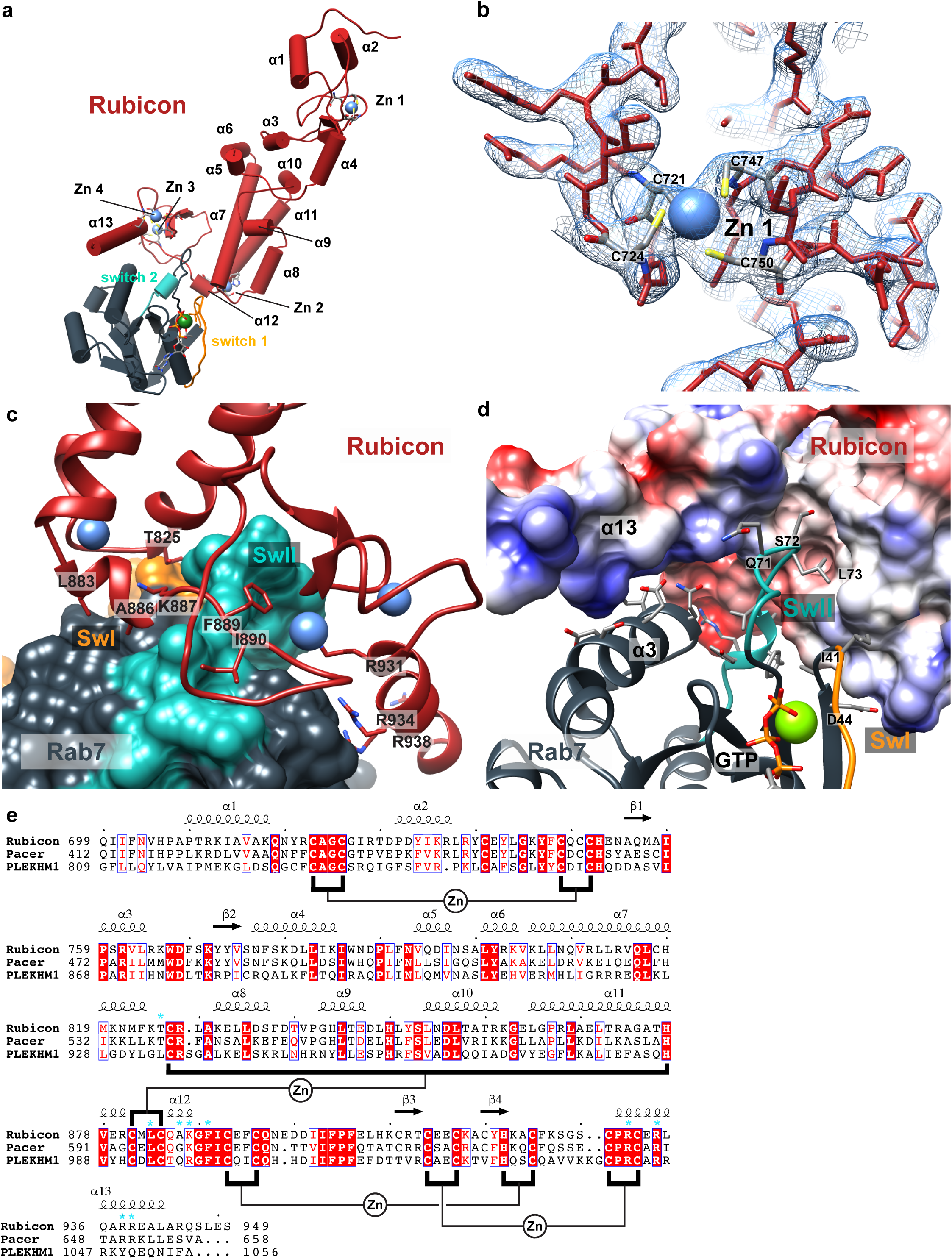
Structure of the Rubicon:Rab7 dimer interface. (a) Cylindrical representation of the Rubicon:Rab7 complex with helices and zinc fingers numbered. Rab7 switch regions are colored and labelled in orange and blue. (b) 2.8Å 2Fo-Fc composite omit map with the final model superimposed. (c) Surface representation of human Rab7 with ribbon representation of the Rubicon RH domain. Key interacting Rubicon residues shown as stick representations. Rab7 SwII shown in blue, and SwI in orange. (d) Surface representation of human Rubicon colored by Coulombic potential and ribbon representation of Rab7 with key residues shown as sticks. (e) Multiple sequence alignment of human Rubicon, Pacer, and PLEKHM1. Secondary structure displayed above the alignment is derived from the Rubicon:Rab7 crystal structure. Zinc finger motifs are annotated in black, indicating which residues are clustered around each divalent zinc atom. Key residues at the Rubicon:Rab7 interface are indicated with blue asterisks. Alignment was generated using ClustalW and ESPript.

The overall structure of Rubicon RH has the shape of the letter ‘L’, where the C-terminal base of the ‘L’ contacts Rab7, and the stalk extends away towards its N-terminus. The ‘L’ is approximately 80 Å long by 50 Å wide. The main RH domain fold consists of four Zn clusters connected to each other by a helical stalk (Fig. 1d, 2a). Zn cluster 1 sits alone at the N-terminal end of the helical stalk. Zn clusters 2-4 form a cluster of clusters at the C-terminal end of the stalk. Zn clusters 1 and 4 are Cys-4 (bound by four cysteines), and Zn 2 and 3 are Cys-2 His-2 (two Cys and two His) clusters (Fig. 1e, f). The latter three of the Zn clusters are close to the Rab7-binding site (Fig. 1f), and the first is distal to Rab7 (Fig. 1e). A pair of helices precedes the RH core, and a single C-terminal helix follows (Fig. 2a).

The overall Rubicon RH fold is unique, as determined by a structural query against the PDB using Dali^19^. Only the region of Zn cluster 1 has any similarity to known structures. The N-terminal part of Rubicon RH (residues 708-784) including zinc cluster 1 resembles the FYVE domain (PDB ID: 1VFY, RMSD = 3.0Å, Fig. 3a-c)^20^. However, the Rubicon Zn cluster 1 lacks some of the basic residues required for canonical FYVE domain phosphatidylinositol 3-phosphate (PI(3)P) binding (Fig. 3c)^20–22^, and Rubicon has not been reported to bind specifically to phospholipids.

**Figure 3.**
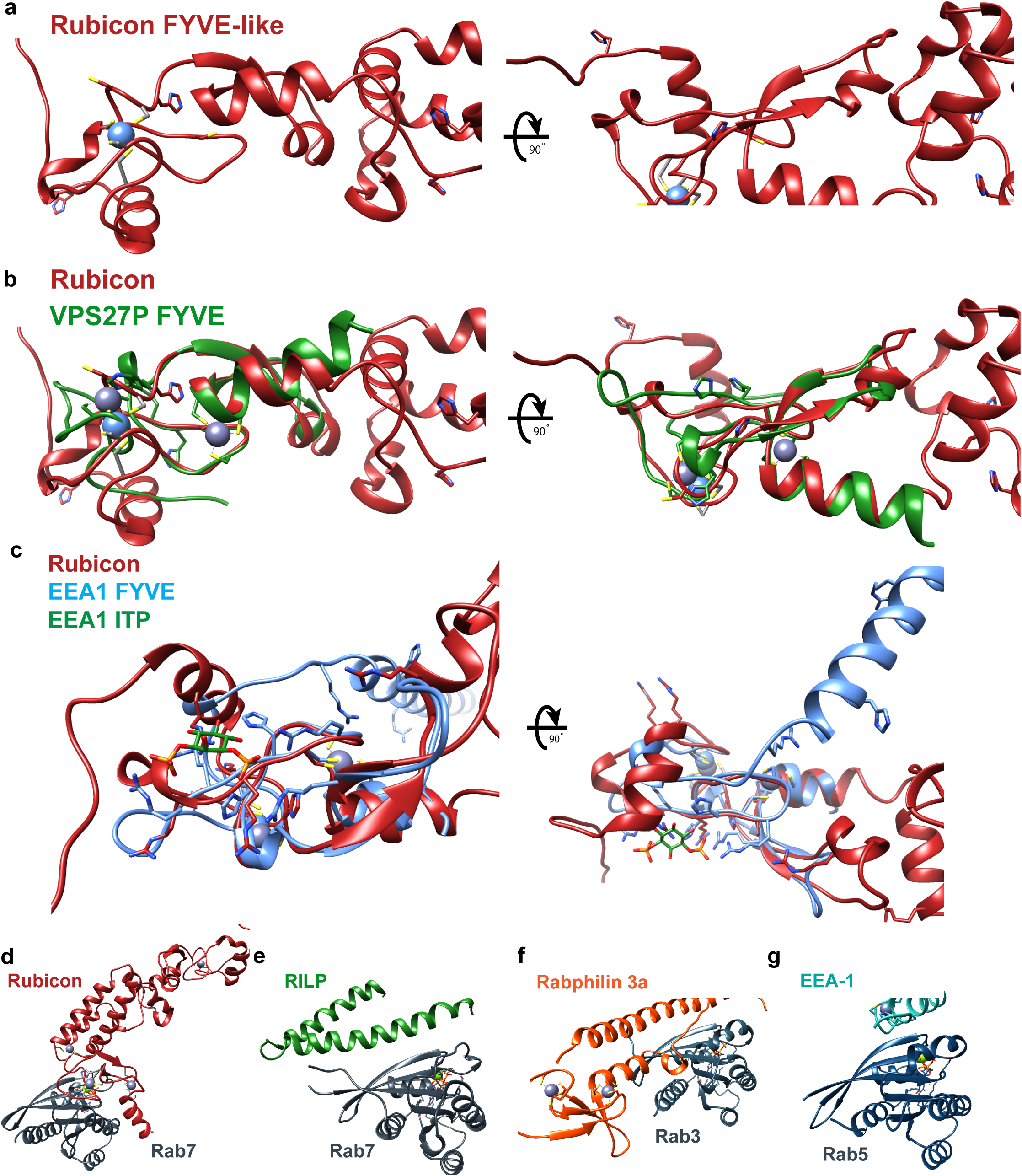
Structural comparison of Rubicon and Rubicon:Rab7 with relevant proteins. (a-c) Comparison of the Rubicon FYVE-like region with FYVE domains of known structure. (a) Cartoon representation of Rubicon FYVE-like region with cysteine and histidine residues shown as sticks. (b) Structural alignment of Rubicon (red) and the FYVE domain of VPS28P (green) (PDB ID: 1VFY). (c) Structural alignment of Rubicon FYVE-like region (red) and FYVE domain of EEA1 (blue) (PDB ID: 1JOC) with key residues and ligands shown as sticks. (d-g) Comparison of the Rubicon:Rab7 complex with other Rab-effector complexes of known structure. (d) Structure of the Rubicon in complex with Rab7. (e) Structure of RILP in complex with Rab7 complex (PDB ID: 1YHN). (f) Structure of Rabphilin 3a in complex with Rab3 (PDB ID: 1ZBD). (g) Structure of EEA-1 in complex with Rab5 (PDB ID: 3MJH).

### The Rubicon RH-Rab7 interface

Rubicon contacts Rab7 via Zn clusters 2-4, the C-terminal end of the main helical stalk, and the C-terminal helix (Fig. 2a, c, d). The interface buries 870 Å^2^ of solvent accessible surface area on Rubicon and 960 Å^2^ on Rab7. Small GTPases contain two principal regions whose conformation changes in response to guanine nucleotide hydrolysis. These regions are known as Switch I and Switch II. At least one of these switch regions is almost always involved in GTP-dependent binding to effectors. Rab7 Sw I is almost completely unencumbered in the Rubicon-bound state (Fig. 2a). In the structure of Rab7 alone, the Sw I and Sw II are both ordered in the presence of the GTP analog GPPNP, while their central elements (residues 35-42 and 65-73) are disordered in the GDP-bound state^23^. In complex with Rubicon RH, we find that Sw I is in a conformation essentially identical to apo Rab7-GPPNP. Sw I Ile41 contacts Rubicon T825, but otherwise Sw I only minimal interacts with Rubicon RH.

Sw II, on the other hand, makes extensive contacts with Rubicon. Sw II is ordered, as seen also in apo Rab7-GPPNP, but its conformation changes considerably due to the many strong interactions with Rubicon RH (backbone r.m.s.d. = 3.1 Å, relative to apo Rab7-GPPNP). The predominantly hydrophobic center of Sw II, residues 72-77, is most extensively involved. Leu73 and Ala76 are major anchors for this segment. Leu73 contacts Rubicon side-chains Met822, Thr825, and the aliphatic parts of Asn821 and Lys887, whilst Ala76 contacts Rubicon Ile890 and Ile900. The Arg79 guanidino group forms a polar interaction network with the side-chains of Rubicon Gln895 and Glu897. Main-chain atoms of Ile73-Gly74 pack against the phenyl ring of Rubicon Phe889. SwII residues 75-78 for a single turn of α-helix. The N-terminal backbone amide groups of this helix donates main-chain hydrogen bonds to C-terminal carbonyl of α12, which emerges from Zn cluster 2. This unusual feature resembles a kinked two-turn helix formed in *trans*.

Several additional non-Switch interactions contribute to the Rubicon RH-Rab7 interaction (Fig. 2c), some with residues immediately C-terminal to Sw I, and some with the highly acidic surface of Rab7 α3. The side chain of Leu883 lies in a hydrophobic pocket formed by Trp62 and the Sw I proximal Phe45 of Rab7. Asp44 and Arg827 form a salt bridge. Multiple Arg residues of the C-terminal helix α13 of Rubicon, notably Arg931, Arg934, and Arg938 interact with Asp100, Asp104, Glu105 of Rab7 α3 and surrounding residues.

Collectively, these interactions drive a unique binding mode as compared to other Rab7 effectors. A structural comparison of Rubicon with selected Rab GTPase:effector complexes is presented in Fig. 3d-g. The binding mode of Rubicon is distinct from those of other Rab7 interactors such as RILP^24^. In addition, while Rab5 interactor EEA1 (Early Endosomal Autoantigen 1)^25^ and Rab3 interactor Rabphilin^26^ both contain zinc finger motifs (Fig. 3f, and g, respectively), neither resembles Rubicon in their Rab interaction geometry.

### The Rubicon:Rab7 structural interface is required for co-localization in cells

To validate the role of key residues at the Rubicon:Rab7 interface *in vivo*, HeLa cells were co-transfected with plasmids encoding mRFP-tagged Rab7 and various FLAG-tagged Rubicon variants (wild type and a number of mutants). Mutants T825D, L883D, A886D, K887D, and F889D introduced negatively charged aspartate residues into the hydrophobic pocket at the SwII-proximal region of the dimer interface. Additional mutants consisted (1) of all the hydrophobic pocket mutation (T825D, L883D, A886D, K887D) together, and (2) four arginine residues on helix α13 of Rubicon substituted for aspartates (R931D, R934D, R938D, R939D). Lastly, the previously-characterized, Rab7 binding-deficient CGHL mutant of Rubicon was used as a negative control. This CGHL mutant has the ZnF cysteines and histidines substituted for glycines and leucines, respectively. Expression of the transfected Rubicon variants was verified by western blotting (Fig. 4a).

**Figure 4.**
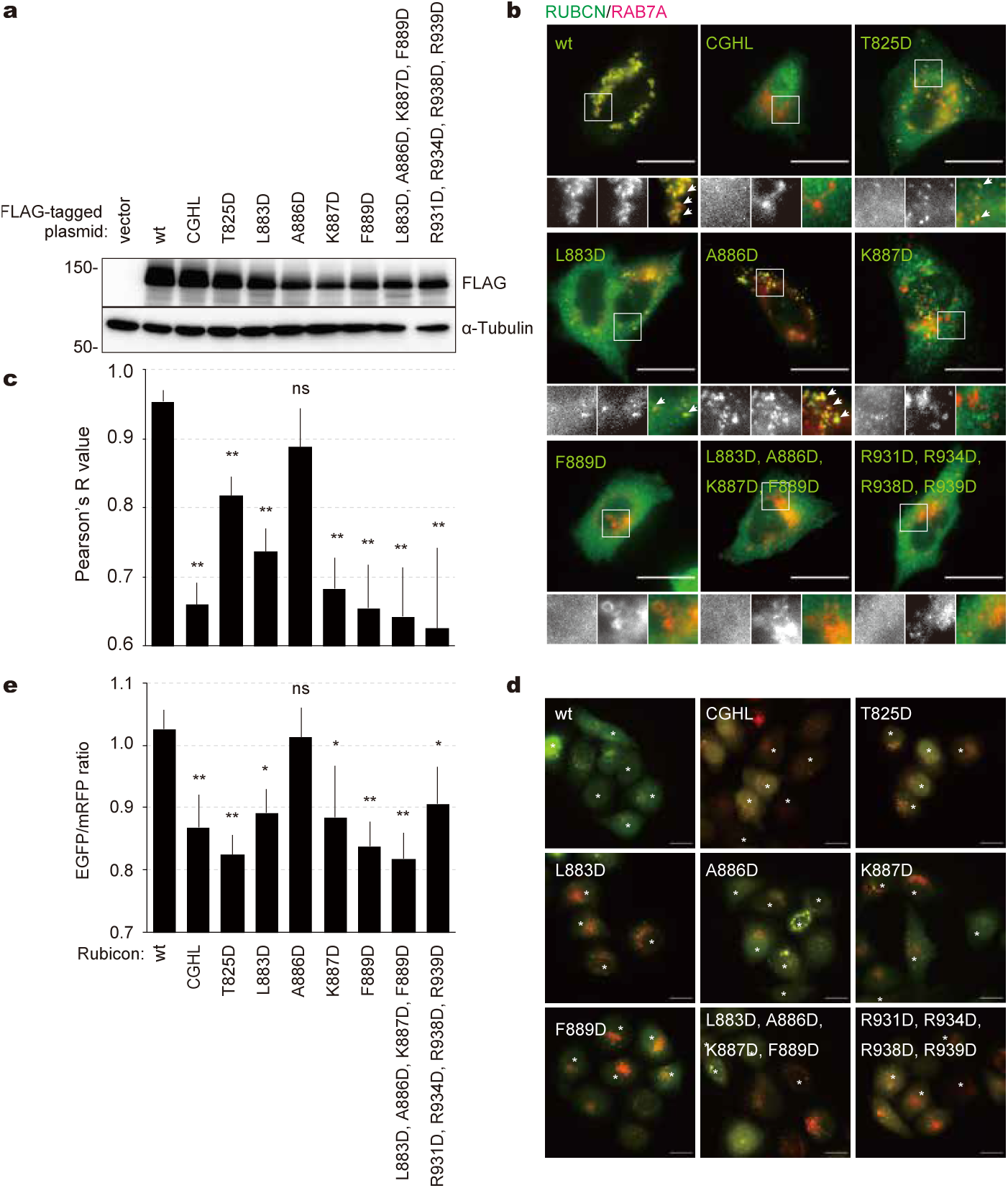
In vivo investigation of the dimer interface. (a) Rubicon mutant expression. HeLa cells were transfected with plasmids encoding 8 FLAG-tagged mutants and wild-type Rubicon and lysed after 24h. Expression levels of Rubicon were detected by western blotting with anti-FLAG antibody. (b and c) Colocalization of Rubicon mutants with Rab7. HeLa cells were co-transfected with FLAG-tagged Rubicon mutants and mRFP-Rab7 and stained with GFP-anti-FLAG antibody after 48h. (b) Rubicon, Rab7, and merged images from boxed areas are magnified and shown from left to right, respectively. White arrows indicate colocalized signals. Scale bars indicate 20μm. (c) Quantification of colocalization between Rubicon mutants and Rab7. Colocalization was analyzed in ImageJ and is shown via Pearson’s R value. Plot show mean values +/- standard deviation (n=3). (d and e) Effect of Rubicon variant overexpression on autophagy. HeLa cells stably expressing tfLC3 (tandem fluorescent LC3; mRFP-EGFP-LC3) were transfected with each FLAG-Rubicon variant plasmid and stained with anti-FLAG antibody. (d) Cells were incubated in starvation medium (S). Cells expressing FLAG-Rubicon are indicated with asterisks. Scale bars indicate 20μm. (e) Signal intensities of EGFP and mRFP were quantified using ImageJ and are shown as EGFP/mRFP ratio normalized to the value from vector-transfected cells. * indicates p < 0.05, ** indicates p < 0.01; ns indicates p > 0.05.

48 hours after transfection, the cells were stained with anti-Flag antibody. As expected, wild-type Rubicon was completely colocalized with Rab7, and the CGHL mutant was dispersed in the cytoplasm and rarely colocalized with Rab7. With the exception of A886D, all of the Rubicon mutations caused a significant reduction in Rubicon-Rab7 colocalization *in vivo* (Fig. 4b,c).

### The Rubicon:Rab7 structural interface is required for autophagy inhibition in cells

To investigate the role of the Rubicon:Rab7 interface in autophagy regulation, we measured autophagic flux in cells overexpressing our Rubicon mutants (Fig. 4d, e). HeLa cells stably expressing tandem fluorescent LC3 (tfLC3, mRFP-EGFP-LC3) were transfected with plasmids encoding the Rubicon mutants. EGFP fluorescence becomes weaker under the acidic conditions of the autolysosome (unlike mRFP), and therefore the EGFP:mRFP fluorescence ratio is expected to be negatively correlated with autophagic flux. As expected, overexpression all Rubicon mutants except for A886D resulted in decreased EGFP:mRFP ratios, indicating increased flux. Wild-type Rubicon overexpression increased the ratio relative to control cells, indicating suppression of autophagic flux.

## Discussion

Rubicon, along with Bcl-2^15^, is one of most prominent negative regulators of autophagy and is therefore considered a promising target for autophagy inducing therapeutics^11, 12^. Despite its significance, however, the only available structural data have been a 6 Å cryo-EM and hydrogen-deuterium exchange characterization of a short portion of the central IDR as bound to PI3KC3-C2^8^. Here, we have provided the first near atomic-resolution information on Rubicon, and the first structure of an RH domain.

The RH domain is the most conserved element of the RH domain-containing family proteins, comprising Rubicon, PLEKHM1, and Pacer. The structure of the Rubicon RH domain can therefore serve as a prototype for the other RH domains. In its RH domain, PLEKHM1 has 32 % sequence homology with Rubicon, and Pacer has 56 % identity. The zinc coordinating residues are identically conserved (Fig. 2e), as is expected given their central role in the structural integrity of the RH domain. The Rubicon residues we identified as biophysically important for Rab7 interaction and autophagy suppression are mostly conserved in PLEKHM1 and Pacer. However, there are some exceptions. The key Ala886 of Rubicon is replaced by Gly and Gln, for example. This suggests that it should be possible to selectively target Rubicon relative to PLEKHM1^9, 27^ and Pacer^17^, which are considered to be positive regulators of autophagy.

The data presented here solidify the significance of the RH-Rab7 interaction in autophagy inhibition by Rubicon, consistent with the finding that disruption of one of the Zn clusters ablates Rubicon function^9^. This finding helps validate that the Rubicon RH-Rab7 interaction is a targetable node in the regulation of autophagy.

Rubicon is a 972-amino acid protein, and the structure and function of most of the rest of Rubicon is still unknown. Apart from the RH domain, and an N-terminal 180-residue RUN domain of unknown function, the remainder of Rubicon consists of intrinsically disordered regions (IDR). Residues 489-638 bind to PI3KC3-C2 and inhibit its ability to bind productively to membranes by blocking the BECN1 BARA domain^8^. Rab5, which is present on early endosomes and autophagosomes, is a positive regulator of PI3KC3-C2^28^. The conversion of Rab5 positive early endosomes to Rab7 positive late endosomes is driven by the Rab7 GEF Mon1-Ccz1^29^. It has been shown, at least in yeast, that Rab7 can also be recruited to, and activated on, autophagosomes by the Mon1-Ccz1 complex^30^. Recruitment of Rubicon, as a negative regulator of PI3KC3-C2, would serve as a termination signal for PI(3)P formation in *cis* upon conversion of endosomes and autophagosomes from Rab5 to Rab7 positive. This simple model is consistent with the literature and the data presented here, but other models are possible. Much still remains to be learned about the structure, function, and physiology of Rubicon, and the data shown here provide an important foundation for progress in this area.

## Methods

### Plasmid construction

Synthetic DNA encoding the Rubicon RH domain (residues 699-949) was synthesized and cloned into a pET vector with an N-terminal His6-MBP tag followed by a tobacco etch virus (TEV) cleavage site. The synthetic DNA sequence was codon-optimized for expression in *E. coli*.

Synthetic DNA encoding human Rab7 (2-183) was cloned into a pET vector with an N-terminal 6xHis tag followed by a TEV cleavage site. The GTP-locking mutation Q67L was introduced using site-directed mutagenesis. All constructs were cloned using the ligation independent cloning (LIC) method.

### Protein expression and purification

All proteins were expressed in *E. coli* BL21 Star (DE3) cells (Agilent Technologies). Cells were grown in lysogeny broth (LB) at 37°C until the OD_600_ was 0.55. The cultures were then chilled in an ice water bath for approximately 15 minutes, induced with 0.35mM isopropyl-β-D-thiogalactoside (IPTG) and incubated with mixing for 18-24 hours at 18°C. Cells expressing Rubicon were grown in media containing 150μm ZnCl_2_.

The cells were pelleted by centrifugation at 4,000g for 20 minutes. The cells were resuspended in lysis buffer containing 50mM Tris-HCl pH 8.0, 300mM NaCl, 10mM tris(2-carboxyethyl)phosphine (TCEP), and 1mM 4-(2-aminoethyl)benzenesulfonyl fluoride hydrochloride (AEBSF). 2mM MgCl_2_ was included in the lysis buffer used for cells expressing Rab7. Cells were lysed by addition of 1mg/mL chicken egg white lysozyme and ultrasonication. The lysates were clarified by centrifugation at 34,500g for 1 hour at 4°C.

The clarified lysates were loaded onto 5mL HisTrap FF Ni-NTA columns (GE Healthcare) equilibrated in 4°C column buffer containing 50mM Tris-HCl pH 8.0, 300mM NaCl, 2mM MgCl_2_, 30mM imidazole, and 5mM TCEP. The columns were washed with 200mL of column buffer, and the tagged protein was eluted with an imidazole concentration gradient from 30mM to 400mM. Fractions were analyzed by SDS-PAGE, and fractions containing pure target protein were pooled. TEV protease was added in a 20:1 mass:mass ratio to cleave affinity tags. Samples were dialyzed against column buffer to reduce imidazole concentration, and cleaved tags were removed by passing the samples over fresh 5mL HisTrap FF columns equilibrated in column buffer.

To measure the ratio of GTP- to GDP-Rab7, a sample of protein was heat denatured to dissociate nucleotide molecules, centrifuged to pellet protein, and the nucleotide-containing supernatant was analyzed using UV spectroscopy coupled to reverse phase HPLC.

Rab7 and Rubicon were pooled in a 1.5:1 molar ratio, and the salt concentration was reduced to 150mM by buffer exchange. The complex was supplied with 2mM GTP and incubated at 4°C for 1 hour prior to purification by size exclusion chromatography on a HiLoad Superdex S75 16/600 column (GE Healthcare) equilibrated in 50mM Tris-HCl pH 8.0, 150mM NaCl, 2mM MgCl_2_, and 5mM TCEP. The elution fractions containing pure Rubicon:Rab7 dimer were determined by SDS-PAGE and pooled.

### Crystallization of the Rubicon:Rab7 Dimer

The purified Rubicon:Rab7 dimer was concentrated to 6mg/mL using a 10kDa molecular weight cutoff centrifugal concentrator (Amicon). During concentration, the buffer was exchanged to a final state of 30mM Tris-HCl pH 7.4, 100mM NaCl, 2mM MgCl_2_, 1mM GTP, and 10mM TCEP. The concentrated sample was centrifuged for 10m at 20,000g at 4°C to pellet precipitated protein, and the supernatant was collected. 200μL of the protein solution was mixed with an equal volume of precipitant solution composed of 100mM Tris-HCl pH 7.4, 6% (w/v) polyethylene glycol (PEG) 8000, and 15mM TCEP. Clusters of crystals were initially obtained, and single crystals were obtained by microseeding with the seed bead method (Hampton Research). Crystals were grown by hanging-drop vapor diffusion at 20°C. Crystals appeared within 5 minutes, and grew to full size overnight. Crystals were cryoprotected using Paratone N (Hampton Research) and flash-frozen in liquid nitrogen.

### Structure Determination

X-ray diffraction data were collected on beamlines 8.3.1 and 5.0.2 at the Advanced Light Source of the Lawrence Berkeley National Laboratory. Data were collected at a wavelength of 1.2830Å. A Dectris Pilatus 3 6M detector (Dectris AG) was used during the data collection. The crystal yielding the best data diffracted to approximately 2.8Å. The reflections were processed using MOSFLM, POINTLESS, AIMLESS, and CTRUNCATE. Statistics for the data collection and processing are provided in Table 1.

An initial, partial molecular replacement solution was obtained using the Rab7 monomer from the Rab7:RILP structure (PDB ID 1YHN) as a search model. However, while the figures of merit from molecular replacement indicated that the solution was correct, the quality of the resulting electron density in the region of Rubicon was insufficient to build a complete atomic model. Given this partial solution with Rab7 placed, we used the recently developed protocol^18^ to build models for Rubicon. With no detectable homologous structures in the PDB, we searched for sequence homologs in UniProt using hhblits^31^, and then used the resulting multiple sequence alignment to predict inter-residue distances and orientations with the deep neural network described in^18^. Network outputs were then converted into restraints, and 3D models were generated by a dedicated Rosetta model building protocol based on restrained minimization. A total of 100 structures that were consistent with the predicted constraints were generated. However, initial placement using Phaser yielded no plausible solutions with TFZ scores above 6. Following visual inspection of the predicted models, we manually trimmed the predicted Rubicon models into three domains, corresponding to residues 699-790 (domain 1), 795-891 (domain 2), and 892-953 (domain 3). Running molecular replacement searches for these domains individually yielded strong hits for domains 1 and 2 (Phaser TFZ values 8.6 and 12.3, respectively), but no hits for domain 3 (TFZ scores below 6). Notably, the termini of the domains were placed in mutually compatible positions suggesting the placement was correct. Following refinement of a model containing Rab7 and domains 1 & 2 of Rubicon with Phenix and Rosetta^32^, and further trimming of domain 3 to 193-242, we were able to identify a good placement of domain 3 (Phaser TFZ score 9.8 and N terminus consistent with domain 2’s placement). Finally, RosettaES^33^, refinement in Phenix/Rosetta^32^, and modelling in Coot^34^ were used to complete the structure.

### Cell Culture and Transfection

HeLa and PlatE cells were grown in Dulbecco’s modified Eagle medium (DMEM) (D6429, Sigma), supplemented with 2mM L-glutamine, 100 U/ml penicillin, 100μg/mL streptomycin, and 10% fetal calf serum. To generate HeLa cells stably expressing tfLC3, HeLa cells were infected with retrovirus generated from pMRX-IRES-puro-tfLC3 plasmid^35^ and selected in medium containing 3μg/ml puromycin. To trigger and visualize starvation-induced autophagy, cells were incubated with Earle’s Balanced Salt Solution (EBSS) for 2h at 37°C. For DNA transfection, we used TransIT-LT1 Transfection Reagent according to the manufacturer’s protocol (Mirus Bio LLC).

### Retrovirus Production

Retrovirus production and cell transductions were performed as described earlier^36^. In brief, PlatE cells were co-transfected with the packaging plasmid pMRX-puro-tfLC3 plasmid and the envelope-encoding plasmid pVSVG by using of polyethylenimine (Polysciences Inc.). Supernatants were harvested 48 h post-transfection and filtered.

### Rubicon Mutant Generation

Rubicon mutants were generated by site-directed mutagenesis and subcloned into pcDNA3.1 plasmids with FLAG sequence.

### Rubicon-Rab7 Colocalization Analysis

HeLa cells were co-transfected with each Rubicon plasmid and mRFP-Rab7^9^ and stained with anti-FLAG M2 antibody (Sigma) at 48h post-transfection.

### Immunofluorescence Microscopy

Cells cultured on glass coverslips were fixed with 4% paraformaldehyde in PBS for 30 min. The cells were permeabilized with PBS containing 0.1% Triton X-100, blocked with 5% FBS, and incubated with diluted anti-FLAG antibody for 60 min at room temperature. After washing with PBS three times, cells were incubated with Alexa405 or Alexa488 labeled secondary antibodies in PBS containing 5% FBS for 60 min. After mounting with VECTASCHIELD (VECTOR Laboratories), images were obtained with an Olympus IX83 microscope.

### Statistics and Reproducibility

Student’s T-tests were performed for paired groups. A p-value less than 0.05 was considered statistically significant. All experiments were repeated two or three times independently, as indicated in the figure legends.

## Data availability

Structural coordinates generated in this work have been deposited in the RCSB Protein Data Bank with accession code 6WCW.

## Acknowledgements

We thank George Meigs of the LBNL Advanced Light Source for advice on crystallographic data collection. We also thank the members of the Hurley Lab for helpful discussions.

## Author Contributions

Conception: H.K.B. and J.H.H.; Methodology: H.K.B., Y.J.I., T.Y., F.D., and J.H.H.; Investigation: H.K.B., K.T., J.M.B., M.H., D.P.F., and I.A.; Writing: H.K.B. and J.H.H.; Review and Editing: all authors; Funding Acquisition: J.H.H., F.D., Y.J.I., and T.Y.; Supervision: J.H.H., F.D., Y.J.I., and T.Y.

